# *Phosphoglycerate mutase 5* regulates lipid metabolism and mitochondrial homeostasis in hepatocellular cancer cells

**DOI:** 10.64898/2026.05.01.718031

**Authors:** Praveen Kumar Guttula, Ganesan Muthusamy, Chin-Chi Liu, Priscilla Devora, Emi Sasaki, Tyler Butsch, Hasand Gandhi, James Moran, Manas Ranjan Gartia, Andrea N. Johnston

## Abstract

The mitochondrial membrane protein phosphoglycerate mutase 5 (PGAM5) is a protein of interest in the complex transition from hepatic steatosis to hepatocellular carcinoma. PGAM5 is a serine/threonine/histidine phosphatase that plays a role in mitochondrial biogenesis, mitophagy, and multiple cell death pathways. Increased expression of PGAM5 in hepatocellular carcinoma is correlated with reduced patient survival. In this study, we demonstrate that loss of PGAM5 alters the bioenergetic landscape of liver cancer by promoting mitochondrial oxidant injury and suppressing the glycerophospholipid and lysophospholipid pathways, leading to accumulation of the bioactive phospholipid lysophosphatidylcholine. Additionally, *PGAM5* deletion downregulates fatty acid biosynthesis, resulting in reduced cellular diacylglycerol concentrations through two probable mechanisms: attenuated long chain fatty acid uptake and suppressed *de novo* synthesis. These findings underscore the broad impact of a single phosphatase on mitochondrial function and provide a rationale for therapeutically targeting PGAM5 to disrupt lipid metabolism in hepatocellular carcinoma.

## Introduction

Metabolic dysfunction-associated steatotic liver disease (MASLD) is a significant risk factor for progression to the inflammatory liver disease metabolic dysfunction-associated steatohepatitis (MASH) and hepatocellular carcinoma (HCC). With the growing obesity epidemic, MASLD/MASH is rapidly becoming a leading cause of HCC. In cancer, unrestrained cellular growth and proliferation impart unique nutrient requirements.^1^ Reprogramming of lipid metabolism is a hallmark of HCC.^2^ When glucose is depleted, fatty acids provide an energetically rich fuel source. Further, lipid droplets serve as depots for membrane components and metabolites necessary to cell signaling.^3^ The majority of HCCs have an increased triacylglycerol content. ^2,4^ The intracellular lipid pool is replenished predominantly by upregulation of *de novo* lipogenesis with enhanced fatty acid uptake as a contributing factor.^5^

Phosphoglycerate mutase 5 (PGAM5) is a protein of interest in the complex transition from metabolic MASLD to HCC.^6–9^ Upregulation of PGAM5 occurs in a subset of patients with HCC, and overexpression correlates with reduced overall survival.^7,10–14^ PGAM5 is an atypical member of the PGAM superfamily. Unlike other family members, which are phosphotransferases or phosphohydrolases of small metabolites, PGAM5 dephosphorylates protein substrates targeting serine, threonine, and histidine residues.^15,16^ PGAM5 contains a WD motif (WDPNWD) essential for multimerization, a catalytic domain that contains residues (H105) required for catalysis, and a C-terminal dimerization domain critical for dimer formation.^17,18^ In states of health, PGAM5 localizes to the mitochondrial intermembrane space where its phosphatase domain is accessible to the cytosol.^19^ In response to mild oxidant stress, PGAM5 promotes mitochondrial biogenesis and mitophagy, but severe oxidant injury accompanied by loss of inner mitochondrial membrane potential leads to PGAM5 cleavage by the intramembrane proteases, presenillin-associated rhomboid- like protein (PARL) and overlapping activity with m-AAA protease (OMA1).^19,20^ This cleavage event releases PGAM5 from its mitochondrial anchor, enabling it to oligomerize into a dimer or dodecamer, which stabilizes its catalytic site.^21^ The functional shift in PGAM5’s location and structure contributes to its diverse cellular effects.^22,23^

Work in our laboratory and by other researchers has demonstrated that reduced PGAM5 expression limits HCC viability.^6,8,10^ These findings have been attributed to increased regulated cell death, deranged mitophagy, disrupted Wnt/β-catenin signaling, and dysregulation of metabolic pathways.^8,20,24–31^ To investigate the physiologic impact of PGAM5 expression on lipid metabolism, we utilized wild type (WT) and *PGAM5* knockout (PGAM5^KO^) HCC and hepatoma cell lines to perform an unbiased assessment of the transcriptome and lipidome. Herein we show that loss of PGAM5 expression in liver cancer cell lines dramatically changes the expression of genes associated with metabolism and the composition of cellular lipid species through suppression of fatty acid uptake and altered mitochondrial bioenergetics. Cumulatively, these findings demonstrate that PGAM5 is a critical co-regulator of metabolism in transformed hepatocytes.

## Materials and Methods

### Canine liver immunohistochemistry

The institutional database was searched for histologically confirmed cases of canine HCC and normal liver samples. A veterinary anatomic pathologist (E.S.) reviewed each case, confirmed the diagnoses, and digitally classified HCC versus peritumoral liver or normal liver.

Seventy de-identified canine samples were evaluated (37 HCC samples, 21 peritumoral samples, and 12 normal liver). Archived formalin-fixed paraffin-embedded (FFPE) liver specimens were sectioned (5 µm) and mounted onto positively charged slides. Formalin-fixed paraffin embedded liver sections (5 µm) were incubated at 55-60°C for 1 hour before deparaffinization and rehydration. Sections were permeabilized in PBS and 0.1% Triton X-100 (Bio-Rad Laboratories) and incubated in antigen retrieval solution (0.1M sodium citrate, pH 6.0) at 95-100°C for 30 minutes with subsequent rinsing in distilled, deionized water. Autofluorescence was quenched by treating sections with TrueBlack (Plus Lipofuscin Autofluorescence Quencher, Biotium). Slides were incubated in 10% normal goat serum for 1 hour at room temperature, washed, and immunolabeled with the primary antibody ([1:100], PGAM5 Rabbit anti-Human Polyclonal IgG antibody, Invitrogen [AB_2900380]) overnight at 4°C. Antibody specificity was confirmed by immunoblot of canine liver tissue. Slides were washed with PBS and fluorescently labeled for 1 hour at room temperature with secondary antibody ([1:400], Alexa Fluor 555-conjugated Goat anti-Rabbit Polyclonal IgG secondary antibody, Invitrogen, [AB_2535849]). Nuclei were stained with Hoechst 33342 Solution [20 µM] (Thermo Fisher) prior to mounting. As per the institutional IACUC committee, use of previously collected, archived tissue samples do not require IACUC approval.

### Imaging and quantitative analysis of protein expression

Fluorescently stained samples were imaged at 20x and 63x magnification (LD Plan-Neofluar 20x/0.4 Corr Ph2 M27, Plan-Apochromat 63x/1.40 Oil M27, Zeiss) using an inverted widefield microscope (Axio Observer Z1, Zeiss) fitted with a xenon arc light source (Lambda DG-4, Sutter Instrument) and reflector cubes (Zeiss 34, 390 nm excitation, 460 nm emission; Chroma 49004, 545 nm excitation, 605 nm emission). Exposures for individual channels (Hoechst 33342, 3000 ms; Alexa Fluor 555, 750 ms) were set by assessing dynamic range (12 bit; 0 – 4096) and signal-to-noise ratio of the images. Channel images were acquired sequentially. Binning was set to 1 × 1 to maximize resolution. Due to the variability in sample staining, the gain was set to one to prevent excessive amplification of fluorescent signals. Images were obtained at 20x magnification (pixel sizes corresponding to 0.323 μm × 0.323 μm) and 63x magnification (0.102 μm × 0.102 μm). Grids were stitched (Zen Blue v. 3.4.91.00000, Zeiss) with a five percent overlap and a maximum shift of ten percent. Stitched grids were saved in a raw, uncompressed format to prevent loss of metadata. Corresponding brightfield images from H&E-stained slides were obtained using a whole slide scanner (Nanozoomer 2.0-HT; Hamamatsu Phototonic) at 40x magnification.

Digital analysis was performed using QuPath© Software (Version 0.4.0). Regions of interest (ROIs) were defined within image grids to exclude vasculature and tissue peripheries for cell detection. ROIs were analyzed for the total number of hepatocytes using QuPath’s built-in cell detection targeting the nuclear stain. Hepatocyte nucleus parameters were adjusted according to area (35 – 95 μm^2^). Intensity parameters were modified to include cells with a mean nuclear intensity ≥ 5. Single measurement classifiers were created to identify hepatocytes and expression of PGAM5. Hepatocyte detection and differentiation from other cell types was based on nuclear morphology (≥ 0.8 circularity, where 1 indicated a perfect circle). The total number of hepatocytes was recorded. Hepatocytes were considered PGAM5-positive if pixels were observed within 5 μm of the nuclei. PGAM5 intensity thresholds for positive pixel detection were iteratively established based on negative antibody controls and pixel intensity of PGAM5 staining. PGAM5 expression per ROI was quantified by counting hepatocytes that contained PGAM5-positive pixels, divided by the total number of hepatocytes. ROIs were averaged to find the mean PGAM5 expression per liver section. A minimum of 5000 hepatocytes were counted per section.

### Cell Lines and Culture Conditions

The Huh-7D 12 (Huh7, 01042712) hepatocellular carcinoma cell line was purchased from Millipore Sigma. Cells were maintained in DMEM, high glucose media (25 mmol/L glucose, Gibco) supplemented with 10% fetal bovine serum (FBS, Avantor, Radnor), and 1% Penicillin–Streptomycin (PS, Gibco) unless otherwise indicated. The HepG2 (HB-8065) hepatoma cell line was purchased from ATCC and maintained in EMEM media (30-2003, ATCC) supplemented with 10% FBS and 1% PS. The Huh7 and HepG2 cell lines were selected because *PGAM5* expression is upregulated compared to normal hepatocytes.^11,12^ As previously reported, the HepG2 PGAM5^KO^ cell line was generated and validated by Synthego Corporation and the Huh-7 PGAM5^KO^ line was generated in our laboratory using PGAM5 CRISPR/Cas9 KO Plasmid (Santa Cruz Biotechnology, Inc., sc-401300).^6^ Nutrient depleted media, composed of DMEM (glucose 12.5 mol/L), FBS 1%, and L-carnitine 0.5 mM (21489, Cayman Chemical), was applied for 16 hours to mimic a reduced macronutrient environment. Cells were treated with bovine serum albumin 80 µM (BSA; 34932, Cayman Chemical) or BSA–Palmitate Saturated Fatty Acid Complex 500 µM (29558, Cayman Chemical) were applied as indicated.

### Measurement of Reactive Oxygen Species (ROS)

The level of intracellular ROS was quantified with the ROS Fluorometric Assay Kit (ThermoFisher, EEA019) as per manufacturer’s instructions. Cells were plated at 5 × 10^5^ cells per well in 12-well plates 24 hours prior to the assay. Fluorescence was measured with a multi-detection microplate reader (Biotek) at 485/530 nm (excitation/emission). Imaging based ROS assessment was collected using a fluorescent microscope (ZEISS Axio Observer Z1). Cells were plated under standard nutrient conditions in 96 well glass bottom imaging plates 24 hours prior to experiments. Nutrient depleted media containing MitoSOX™ Green FM (Invitrogen, M7514, 490/516 nm, 50 nM), MitoSOX™ red superoxide indicator (Invitrogen, M36008, 396/610 nm, 100nM), and LysoTracker Blue DND-22 (Invitrogen, L7525, 373/422 nm, 50 nM) was replaced and cells were incubated for 30 minutes at 37°C with 5% CO2. Immediately prior to imaging, cells were gently washed three times with HBSS (Gibco, 14025092) and imaged.

### Sample preparation, processing, RNA sequencing, and bioinformatic analysis

Cells were trypsinized and washed with PBS. Cell viability and count were performed manually, and the concentration was adjusted to approximately 10,000 cells per sample to optimize input into the Chromium Controller (10x Genomics), per manufacturer guidelines. The Chromium Next GEM Single Cell 3’ Gel Bead Kit v 3.1 (10x Genomics) was used to create Gel Beads-in-emulsion (GEMs), which allowed for the labelling of individual cells’ poly-A mRNA with a 16nt 10x Barcode and a 12nt unique molecular identifier (UMI) as per manufacturer’s instructions. GEMs were lysed and cleaned using DynaBeads (MyOne SILANE).

Oligonucleotides underwent reverse transcription to create barcoded, full-length cDNA from poly-A mRNA. Ten to 12 PCR cycles were performed on a BioRad C1000 Thermal Cycler to amplify the barcoded cDNA. Beckman Coulter SPRIselect Reagent and magnetic Separator was used to clean barcoded cDNA. Total cDNA quality and quantity were assessed using the Agilent Fragment Analyzer. As per the protocol, 25% of total cDNA was used for library preparation.

Sequencing libraries were assessed by the Agilent Fragment Analyzer for quality and quantity. Samples were pooled in equimolar ratios. The Illumina NextSeq 500 was used for library sequencing using a High Output Kit v2.5 (150 cycles). According to 10X Genomics guidelines, Illumina reads were set to 28 × 8 × 0 × 91. RNA sequencing was performed by GeneLab at Louisiana State University.

The cellranger mkfastq and cellranger count tools from 10X genomics, with default settings, were used to process raw sequencing data. The cellranger count was mapped to the GRCh38 human reference provided by 10X genomics. Raw data processing was done separately for each sample. After processing raw sequencing data, the R package Seurat was used to perform all remaining analyses. Data was filtered by housekeeping genes (*ACTB* and *GAPDH*) and mitochondrial genes. The SCTransform function in Seurat was used to normalize the data across samples within the dataset. Enrichment analysis of differentially expressed genes (DEG) were analyzed using the Gene Ontology Resource. Predicted protein interaction analysis was conducting using the String database (version 12.0). Bioinformatics were performed at University of Arkansas College of Medical Sciences.

### Quantitative PCR

Total RNA was extracted from cultured cells using the RNeasy kit (Qiagen) according to manufacturer’s instructions and quantified using a Nanodrop™ (Thermo Fisher). Reverse transcription and quantitative (q) PCR were performed per manufacturer’s protocols using qScript™ cDNA SuperMix and PerfeCTaSYBR® Green FastMix® Reaction Mix (QuantaBio). The housekeeping genes 18s and ß-Actin were used for standardization of sample mRNA content. Analysis was performed on 7900 HT Applied BioSystems™ Real-Time PCR Detection System (BioRad). Each quantitative RT-PCR was performed with 12.5 ng cDNA. The relative expression levels of the target genes were calculated and expressed as -ddCt relative to 18S RNA expression in WT cells. The primers were purchased from Integrated Data Technologies (IDT); sequences are listed below.

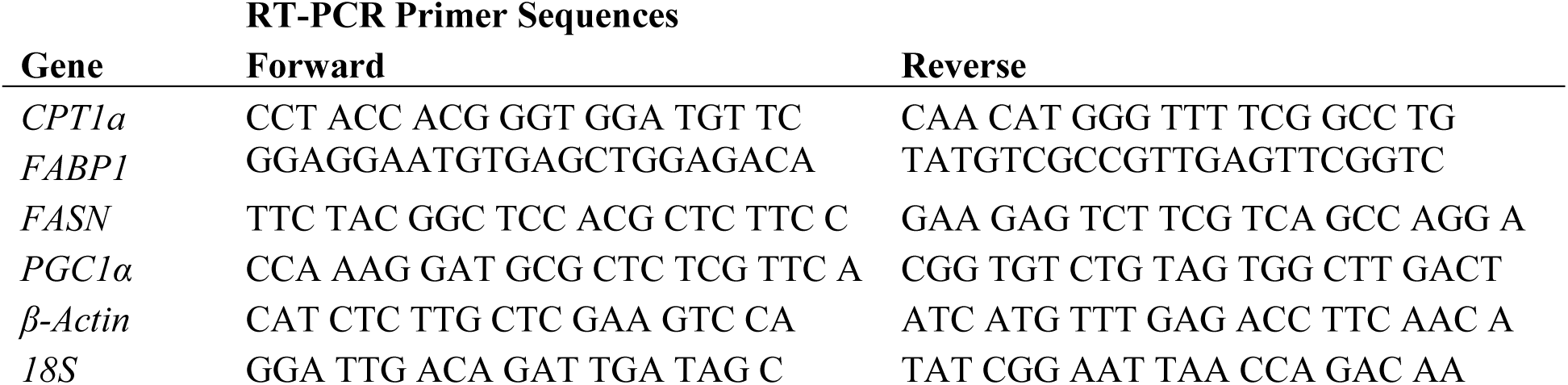

### DAG fluorometric assay

Diacylglycerol concentrations were measured using an enzyme-linked fluorescence assay according to kit protocols (Cell Biolabs, MET-5028). Fluorescence product was measured using 530-nm excitation and 590-nm emission. HepG2 and Huh7 cells were treated overnight with BSA or BSA-conjugated palmitate as previously described.^6^ The media was removed, and cells were washed with ice cold PBS prior to sonication in methanol, followed by extraction of organic phase in chloroform. Total protein was measured by the BCA Protein Assay Kit (ThermoFisher, 23227) and used to normalize fluorescence measurements.

### Gel electrophoresis and Immunoblots

Total cell protein lysates were prepared as previously described.^32^ The protein content was determined using a BCA Protein Assay Kit (ThermoFisher, 23227). Total protein (20 - 40 μg) was separated by SDS-PAGE and transferred onto PVDF membranes. Membranes were incubated with primary antibodies at 4°C overnight and subsequently incubated with their corresponding HRP-labeled secondary antibodies. Bands were detected using ECL (Cytvia, RPN2106) on myECL Imager (ThermoFisher, G2236X). The protein densitometry was measured using ImageJ software (National Institutes of Health). Primary antibodies included: Actin (Invitrogen, AB_2223496), Acetyl-CoA Carboxylase (Cell Signaling, AB_2219400), FASN (Invitrogen-AB_11156419), Lipin-1 (Cell Signaling Technology, 5195, AB_10694491), PGAM5 (Invitrogen, AB_2900380), and PGAM5 (Invitrogen, AB_2900380; canine tissue). Secondary antibodies included: Goat anti-Rabbit HRP (Invitrogen, AB_228341), Anti-mouse (Invitrogen, AB_228295). Lipin-1 was immunoprecipitated from WT and PGAM5^KO^ Huh7 cell lysates using Dynabeads conjugated to lipin-1 antibody (Cell Signaling Technology, 5195). Purity of immunoprecipitated protein was confirmed by immunoblot. Immunoprecipitated protein was subjected to electrophoresis using a 12.5 % SuperSep™ Phos-tag™ (FUJIFILM Wako Pure Chemical Corporation) as per manufacturer’s instructions. The gel was stained with Coomassie Brilliant Blue R250.

### Lipid extraction

Huh7 cell lines were plated at 1 × 10^6^ cells per well in triplicate and cultured under standard conditions for 36 hours. At 36 hours, cells were washed with PBS and detached with trypsin EDTA 0.1%. Following centrifugation, cells were washed thrice with PBS and snap frozen in liquid nitrogen prior to storage at -80°C. Samples and reagents used for extraction were kept on ice throughout the procedure. Cells were lysed in methanol containing 0.1% butylated hydroxytoluene. Cells were vortexed and sonicated for a total of 15 minutes. Cell lysate was cleared via ultracentrifugation at 15,000 g. 190 μL of supernatant was transferred to a new tube and 100 μL of H2O and 500 μL methyl-tert-butyl ether (MTBE) were added. The mixture was vortexed for 30 seconds and incubated on ice for 30 minutes, followed by centrifugation. After centrifugation, 400 μL of the MTBE phase was transferred to a glass vial, 50 μL of H2O was added, and the mixture was briefly vortexed. The extract was dried under a gentle stream of nitrogen and then resuspended in 50 μL of isopropanol.

### Lipidomics and bioinformatics

LC-MS was used to identify lipid species and abundance using a Waters Synapt XS mass spectrometer. All samples were diluted 1:5 and a pool QC was created by combining equal volumes of each sample. A volume equal to 2 µL in positive mode and 4 µL in negative mode was injected in the system. The column used was a Waters Acquity Premiere CSH C18 (100×2.7, 1.7 µm particles and 130 Å pore size) with VanGuard Fit guard column. The mobiles phases were A= H2O:ACN, 4:6 with 0.1 formic acid and 10 mM ammonium formate, and B = IPA:ACN, 9:1 with 0.1% formic acid and 10 mM ammonium formate. Samples were loaded at 40% B. A gradient program was applied as follows: 0-2 min 40% B to 43%B, 2 min to 2.1 min 43% B o 50% B, 2.1 min to 12 min 50% B to 54% B, 12 min to 12.1 min 54% B to 70% B, 12.1 min to 18 min 70% B to 99% B, 18 min to 18.1 min to 40% B, re-equilibration for 1.9 min. The mass spectrometer was operated in “HDMSE” mode, with alternating low and high fragmentation energy spectra and ion mobility active. The acquisition rate was set at 0.3 s, and the collision energy was ramped from 10 to 40 V during the high energy scan. The emitter voltage was set at 1.5 kV in positive mode and 2 kV in negative mode. Data were manually checked with MassLynx 4.2 and then used for automated data analysis using Progenesy QI (ver. 3.0). Global sample analysis was performed and pathway analysis was completed using BioPAN (https://www.lipidmaps.org/biopan/; Z > 1.645 corresponds to p < 0.05).

### Seahorse assays

Huh7 cells were plated under standard culture conditions 24 hours prior to metabolic assays in 96 well plates at a density of 40,000 cells per well. Seahorse XF MitoStress Test and Seahorse XF Mito Fuel Flex Test (Agilent) were performed according to manufacturer’s instructions.^33^ The Seahorse XF Pro analyzer (Agilent) was used to acquire data and analyses were performed using Agilent Seahorse Analytics software. Cell quantification was performed after each assay via Hoechst nuclear stain count using the Cytation 5 (BioTek) and label free cell confluence was determined using the Incucyte (Sartorius) bright field analysis.

### Acetyl-CoA assay

Acetyl-CoA concentration was measured using the CheKine™ Micro Acetyl Coenzyme A Assay Kit (MyBioSource, MBS9719208) per the manufacturer’s protocol. In brief, cells were incubated overnight in nutrient replete media. Wells were washed thrice with phosphate buffered saline prior to cell collection and centrifugation. Cell pellets were lysed, and protein concentration was determined using a BCA assay (Thermo Fisher). Samples were incubated with working solution for 10 minutes at 37°C and absorbance (340 nm) was measured at 20 seconds and 1 minute and 20 seconds. Acetyl-CoA concentration was calculated based on the standard NADH curve and protein concentration.

### Transmission electron microscopy

Samples for transmission electron microscopy were fixed in 2.5% glutaraldehyde and 2.5% paraformaldehyde in 0.1 M cacodylate buffer at 4°C for 24 hours. Following fixation, samples were treated with 1% osmium tetroxide and dehydrated in a graded acetone series. Samples were infiltrated and embedded in Poly/Bed 812 resin (Polysciences, Warrington, PA). Thin sections (70 nm) were obtained with a PTXL ultramicrotome (RMC, Boeckeler Instruments, Tucson, AZ, USA) on 200 mesh copper grids, and stained with uranyl acetate and lead citrate. Samples were subsequently washed in cacodylate buffer and routinely processed for electron microscopy. Sections were imaged using a JEOL 100CX Transmission Electron Microscope (Japan) at a 100 kV accelerating voltage.

### Isotope-Ratio Mass Spectrometry (IRMS) measurement of ^13^C-palmitate uptake

The Huh7 WT and PGAM5KO cell lines were treated with 500 uM ^13^C-palmitate (Cambridge Isotope Labs, CLM-8390-0.25) conjugated to fatty acid free BSA or BSA control in DMEM low glucose (1 g/L) media supplemented with 1% FBS for 0, 4, and 16 hours. Duplicate samples were generated for each time point. Cell pellet and supernatant samples were individually loaded into pre-weighed tin capsules (Elementar, 200014577), and dried overnight at 50°C, and then re-weighed to determine sample mass. Quantitation of ^13^C isotope was completed using an Elementar Vario Isotope Cube elemental analyzer (Elementar Analysensysteme GmbH, Hanau, Germany) interfaced to an Isoprime VisION Isotope Ratio Mass Spectrometer (IRMS, Elementar UK Ltd, Cheadle, UK). Samples were combusted at 950°C with excess oxygen in a quartz reactor packed with copper oxide and silver wool. Sample gases were passed through a reduction furnace (650°C) packed with reduced copper. After reduction, the sample gases were passed through a water trap (Elementar, USA) followed by an adsorption trap (40°C) which selectively retains the CO2. The adsorption trap was heated (150°C) to release the CO2 to the IRMS for measurement. Several replicates of at-least two different laboratory reference (previously calibrated against international reference standards) materials were analyzed. The raw isotope values of the samples were calibrated using the standard runs and a two-point calibration. Final abundance of ^13^C was reported as atom percentage (%) excess.

### Statistical Analyses

Statistical analyses were performed using JMP Pro 18.2.0 (JMP Statistical Discovery LLC, Cary, NC). Graphs were created in GraphPad Prism 10.0.2 for Windows (GraphPad Software, LLC, San Diego, CA). Data are expressed as model estimated least-squares mean (LS means) ± standard error of the mean (SEM), unless otherwise noted. For cell culture experiments, each independent experiment performed on a separate day was considered a biological replicate. Technical replicates represent repeated measurements within the same experiment. Technical replicates were averaged prior to statistical analysis, and biological replicates were used as the experimental unit for all statistical tests. For experiments involving genotype and treatment conditions, data were analyzed using analysis of variance (ANOVA) models including genotype, treatment, and their interaction as fixed effects. Because experiments were repeated across independent experimental runs, experiment was included as a random effect in mixed-effects ANOVA models to account for between-experiment variability. When a significant overall effect was detected, pairwise comparisons between groups were performed using Tukey’s HSD post hoc test to adjust for multiple comparisons. Comparisons between two groups were performed using Student’s t-tests. For protein expression analyses, one-tailed t-tests within cell lines were used based on a priori directional hypotheses that deletion of PGAM5 was expected to reduce expression of lipid metabolism–related proteins. For the IR-MS data, mixed-effects ANOVA models were fitted with genotype, time, substrate, sample type, and their interactions specified as fixed effects; experimental batch was included as a random effect. When a significant fixed effect was detected, post hoc comparisons were conducted using Tukey’s HSD tests. Model assumptions for all parametric models were assessed via standardized residual plots and normal Q-Q plots; residuals met assumptions of normality and homoscedisticity. Sample sizes were based on prior similar studies and standard practices in the field. Statistical significance was set at P < 0.05.

## Results

### PGAM5 is overexpressed in hepatocellular carcinoma

Overexpression of PGAM5 is reported in human breast, gastric, lung, skin, colorectal, and liver cancer and PGAM5 has been associated with enhanced mitochondrial quality control and metabolic adaption.^34–37^. To determine if this feature was conserved in spontaneously occurring hepatocellular neoplasms of a non-human species, PGAM5 expression was quantified in the livers of dogs with and without HCC. As in human HCC, immunohistochemical (IHC) analysis (**Figure 1a**) revealed that PGAM5 expression was significantly higher in HCC tissue than peritumoral tissue or normal liver (**Figure 1b**). PGAM5 protein expression in peritumoral and normal liver regions was not significantly different. These results establish the biological relevance of PGAM5 in HCC across species.

**Figure 1.**
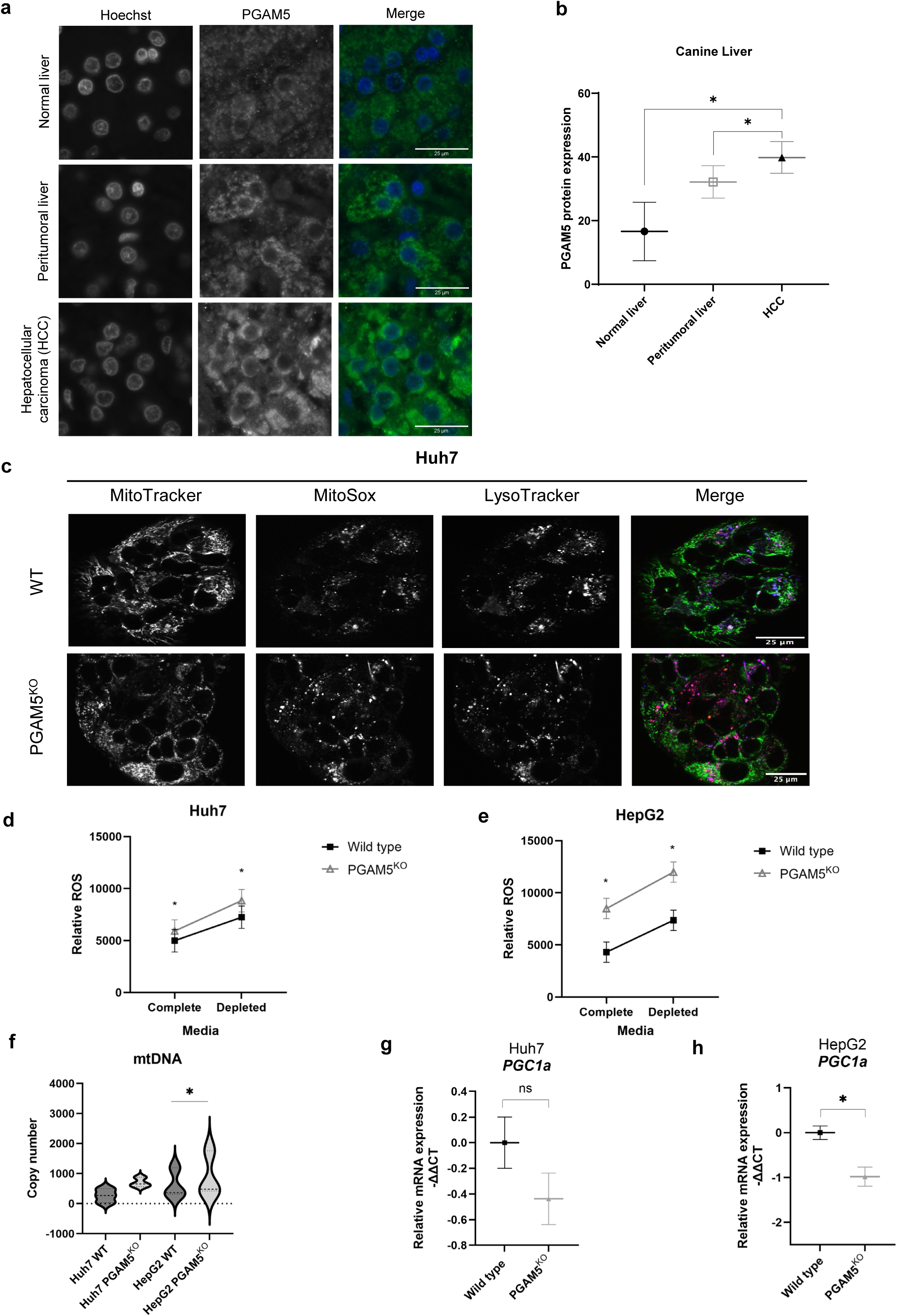
PGAM5 expression mitigates oxidant injury in hepatocellular neoplasms. **a**. Representative images of PGAM5 (green) immunohistochemical labeling in canine normal liver, peritumoral liver, and HCC. Nuclei are stained with Hoechst (blue). **b**. Quantification of immunohistochemical PGAM5 expression in dogs; data include normal liver (n = 12), peritumoral liver (n = 21), and HCC (n = 37). Data expressed as mean ± SEM of percentage of PGAM5-positive cells. **c.** Images of mitochondria (MitoTracker, GreenFM), superoxide (MitoSOX, Red), and acidic lysosomes (LysoTracker, Blue) in WT and PGAM5^KO^ Huh7 cells 30-60 minutes after exposure to nutrient depleted media. **d. e.** Fluorometric assay measurement of ROS in WT and PGAM5^KO^ cells after 16-hour exposure to complete or depleted culture media in Huh7 (d) and HepG2 (e) cells. Data expressed as mean ± SEM. **f.** Violin plots of mitochondrial DNA (mtDNA) copy number in WT and PGAM5^KO^ Huh7 and HepG2 cells. **g., h.** Real-time (RT) qPCR of Huh7 (g.) and HepG2 (h.) wild type (black square) and PGAM5 ^KO^ (grey triangle) *PGC1a* transcripts. Data represent 3 biological replicates with a minimum of 2 technical replicates per sample. Data expressed as mean ± SEM; *, p<0.05.

### Increased ROS production and impaired mitochondrial clearance secondary to PGAM5 deletion leads to persistence of damaged mitochondria

PGAM5 inhibits ROS-induced oxidative stress in some models.^38,39^ To test if deletion of *PGAM5* counteracts this protective effect in HCC, ROS was quantified in WT and PGAM5^KO^ cells. PGAM5^KO^ cells demonstrate increased mitochondrial superoxide radical levels and colocalization of damaged mitochondria with lysosomes (**Figure 1c**). Under nutrient replete conditions, total cellular ROS was significantly greater in the PGAM5^KO^ HCC and hepatoma cells compared to WT cells and this was aggravated by limiting nutrient supply (**Figure 1d, e**). Increased superoxide levels stimulate mitophagy to eliminate oxidant damaged organelles.^40,41^ PGAM5 is known to regulate KEAP1 and Parkin mediated mitophagy.^42–44^ To evaluate whether damaged mitochondria were effectively cleared in PGAM5^KO^ cells, mitochondrial DNA (mtDNA) copy number (**Figures 1f**). An increase in mtDNA was identified in both PGAM5^KO^ cell lines but was only significantly upregulated in HepG2 cells. The relative mRNA expression of *PPARGC1A (PGC1α)* was quantified to assess whether the increased mtDNA resulted from compensatory mitochondrial biogenesis rather than failed mitophagy (**Figures 1f-h**). The lack of *PGC1α* upregulation suggests that energetically compromised mitochondria persist in PGAM5^KO^ cells, consistent with observations in retinal pigment epithelial cells.^45^

### PGAM5 deficiency leads to impaired mitochondrial electron transport and altered nutrient dependency

To assess the impact of ROS accumulation on cellular metabolism, the mitochondrial oxygen consumption rate (OCR) and extracellular acidification rate (ECAR) were quantified in Huh7 WT and PGAM5^KO^ cells. The OCR was significantly downregulated in Huh7 PGAM5^KO^ cells during basal and maximal respiration measured after carbonyl cyanide-p-(trifluoromethoxy) phenylhydrazone (FCCP) mediated uncoupling of oxidative phosphorylation (**Figure 2a**). These results indicate that electron transport chain capacity is impaired upstream of ATP synthase, consistent with dysfunction at complex I and/or III. The ECAR, reflective of glycolysis, was also reduced in the Huh7 PGAM5^KO^ cells compared to WT cells at all time points (**Figure 2b**). Taken together, these data suggest a metabolically quiescent phenotype in PGAM5^KO^ cells. We next asked if specific macronutrient metabolic pathways were impacted by *PGAM5* deletion in the Huh7 cell line. The mitochondrial fuel flexibility assay was performed to test if PGAM5^KO^ HCC cells utilize fuel sources (fatty acids, glucose, and glutamine) differently than WT HCC (**Figure 2c**). The WT cells demonstrated high basal utilization of fatty acids and glucose (**Figure 2d**).

**Figure 2.**
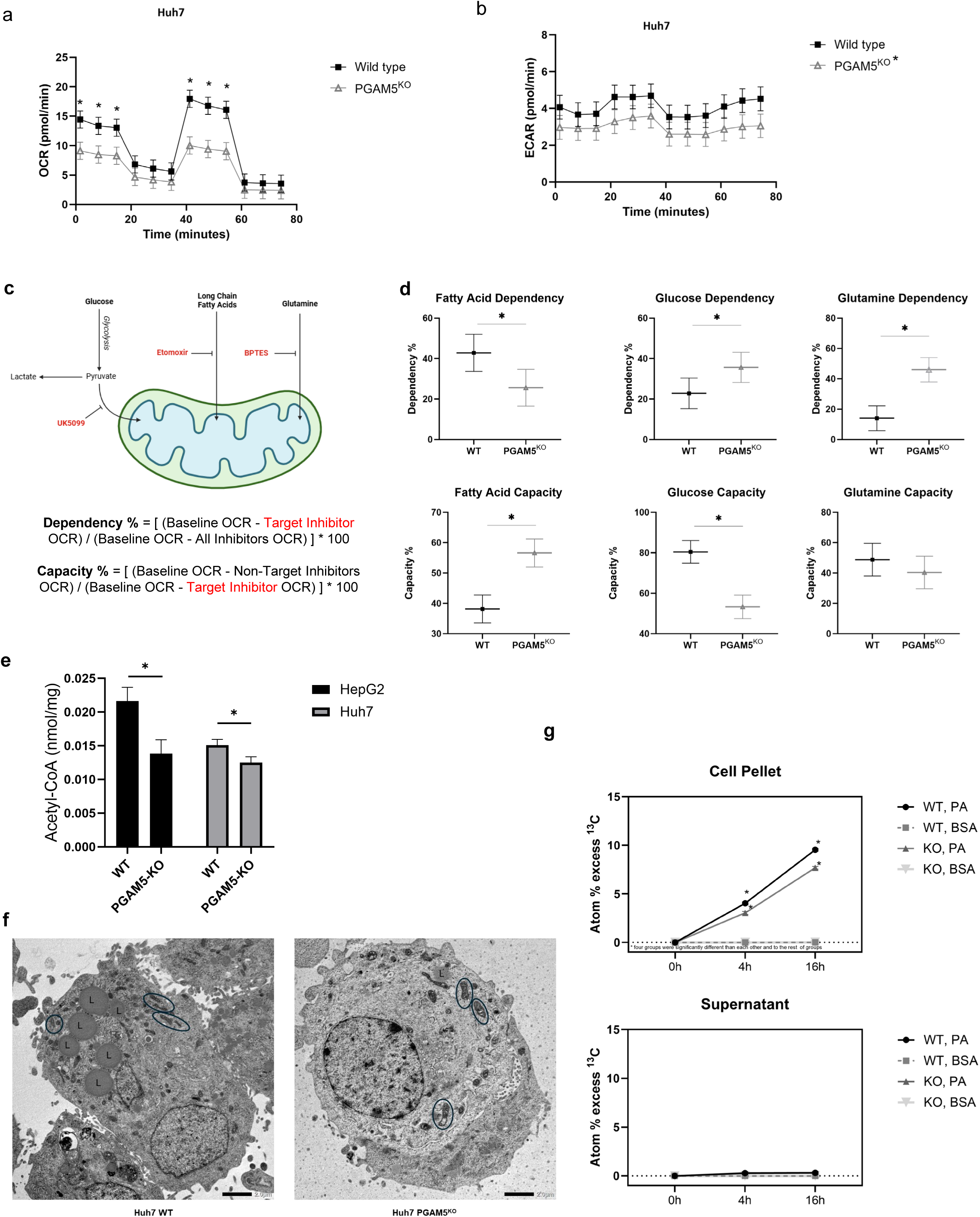
Deletion of *PGAM5* dysregulates mitochondrial homeostasis. **a**. Graph of oxygen consumption rate (OCR) in wild type (WT, black squares) and *PGAM5* knockout (PGAM5^KO^; grey triangles) Huh7 cells. Displayed data are derived from 3 biological replicates with a minimum of 3 technical replicates per sample. Results are normalized to cell count. **b**. Graph of extracellular acidification rate (ECAR) in WT (black squares) and PGAM5^KO^ (grey triangles) Huh7 cells. **c.** Schematic of the Agilent Seahorse fatty acid fuel flex assay. **d.** Dependency percentage and capacity percentage of fatty acids, glucose, and glutamine in WT (black circles) and PGAM5^KO^ (grey triangles) Huh7 cells. Displayed fuel flex data are derived from 3 biological replicates with a minimum of 3 technical replicates per sample. **e.** Acetyl-CoA concentration in WT and PGAM5KO HepG2 and Huh7 cells. Data represent 3 biological replicates with a minimum of 2 technical replicates per sample **f.** Representative transmission electron micrographs of Huh7 WT and PGAM5^KO^ treated with 500 uM palmitate for 16 hours. Black ovals encircle mitochondria, “L” labels lipid droplets, and scale bars equal 2.0 µm. **g.** Isotope-Ratio Mass Spectrometry atom percentage (%) excess of ^13^C in Huh7 WT and PGAM5^KO^ pelleted cells or supernatant following treatment with 500uM ^13^C-palmitate (PA) or unlabeled bovine serum albumin (BSA) for 16 hours. Data represent 3 biological replicates with 2 technical replicates per sample; *, p<0.05.

While dependence on glucose and glutamine was significantly increased after *PGAM5* knockout, basal metabolism of fatty acids was significantly decreased suggesting a preferential reliance on glutamine anaplerosis and glucose oxidation rather than beta-oxidation. Paradoxically, when glucose and glutamine metabolism were chemically inhibited, PGAM5^KO^ cells significantly upregulated fatty acid oxidation indicating that the latent capacity to utilize fatty acids was maintained despite mitochondrial oxidant injury. To better characterize the net effect of metabolic flux caused by *PGAM5* deletion, we quantified total cellular acetyl-CoA concentration under nutrient replete conditions. The metabolite acetyl-CoA can be derived from amino acid, fatty acid, and glucose catabolism.^46,47^ In fed states, acetyl-CoA is used for *de novo* lipogenesis, but during fasted states, acetyl-CoA is oxidized in the (tricarboxylic acid) TCA cycle for ATP synthesis.^46,47^ In both cell lines, acetyl-CoA concentration was significantly reduced in PGAM5^KO^ compared to WT cells (**Figure 2e**).

To clarify if the reduced level of acetyl-CoA in PGAM5^KO^ cells reflected diminished fatty acid oxidation due to mitochondrial dysregulation versus a decrease in fatty acid concentration, transmission electron microscopy was used to assess lipid droplet accumulation following treatment with palmitate. Electron micrographs documented the previously reported mitochondrial morphologic changes associated with *PGAM5* knockout (**Figure 2f**).^45,48,49^ Further, PGAM5^KO^ cells appeared to have fewer lipid droplets than WT cells and demonstrated significantly reduced ^13^C-palmitate uptake (**Figure 2g**). Taken together, these data suggest that *PGAM5* deletion causes mitochondrial bioenergetic dysregulation, attenuating mitochondrial metabolic activity and leading to distinct changes in nutrient uptake and utilization.

### PGAM5 modulates gene expression of metabolic regulators

We hypothesized that PGAM5 could modulate mitochondrial-nuclear crosstalk to direct nutrient uptake and allocation. To assess the transcriptional regulatory pattern of PGAM5 in HepG2 and Huh7 cells, we performed RNA sequencing (RNAseq) on WT and PGAM5^KO^ cells. Amongst the top 20 genes, four nuclear encoded transcripts associated with the electron transport cycle were downregulated: *ATP5MC1* (*ATP Synthase Membrane Subunit C Locus 1,* complex V), *ATP5ME* (*ATP Synthase Membrane Subunit E,* complex V), *NDUFS6* (*NADH: Ubiquinone Oxidoreductase Subunit S6,* complex I), and *NDUFB2* (*NADH: Ubiquinone Oxidoreductase Subunit B2I,* complex I, **Figure 3a**).^50^ Gene ontology analysis of the Huh7 RNAseq dataset identified enrichment in metabolic and mitochondrial pathways across the biological processes, cellular components, and molecular functions ontology groups (**Figure 3b**). A STRING analyses of the Huh7 and HepG2 RNAseq data demonstrated that lipid metabolism related transcripts were significantly downregulated in PGAM5^KO^ cells (**Figure 3c** (Huh7); **Figure 3d** (HepG2)).

**Figure 3.**
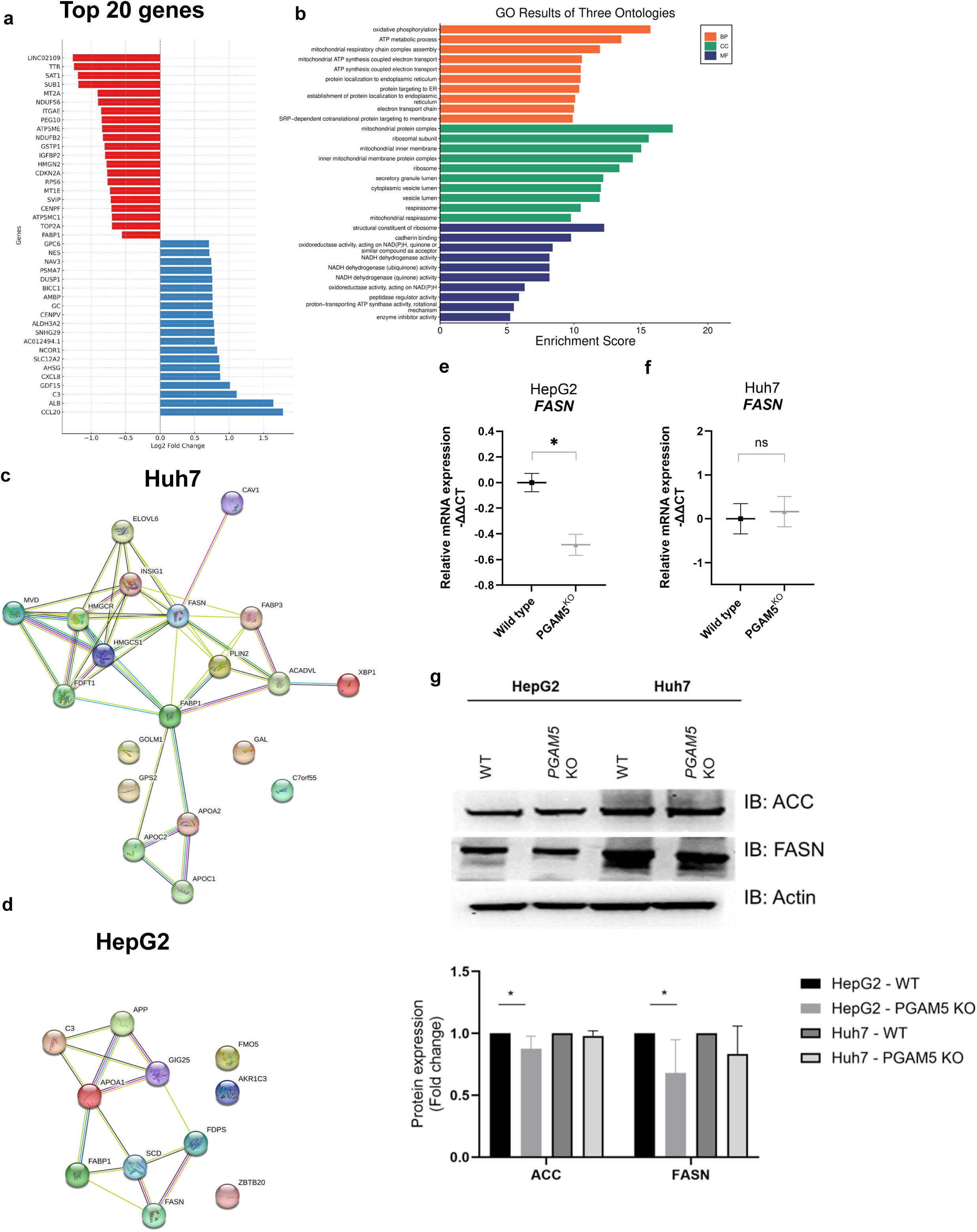
RNA sequencing data link *PGAM5* expression to metabolic pathways. **a.** Top 20 downregulated genes (red) and upregulated (Blue) genes secondary to *PGAM5* deletion in Huh7 cells. **b**. Gene ontologies (GO) plot of enriched biological processes (BP, orange), cellular components (CC, green), and molecular functions (MF, blue) associated with PGAM5 expression in Huh7 cells. **c., d.** STRING analyses of Huh7 (**c**) and (**d**) HepG2 GO regulation of lipid metabolism (GO:0019216). PPI enrichment p-value 1.06e-10, FDR < 0.02. **e., f.** Real-time (RT) qPCR of HepG2 (**e**) and Huh7 (**f**) wild type (black square) and PGAM5 ^KO^ (grey triangle) of *fatty acid synthase* (*FASN*). Data are derived from a minimum of 2 biological replicates with 2 technical replicates per sample and are represented as mean ± SEM; *, p<0.05. **g.** Representative immunoblots of acetyl-CoA carboxylase (ACC), FASN, and actin from WT and PGAM5^KO^ HepG2 and Huh7 cell lysates. Relative protein expression is normalized to actin. Data represent 3 biological replicates presented as mean ± SD; *, p<0.05.

Notably, both *fatty acid binding protein (FABP1)*, a long chain fatty acid importer, and *fatty acid synthase (FASN)*, a critical catalyst in *de novo* lipogenesis, were significantly downregulated in both PGAM5^KO^ cell lines. We have previously shown that *FABP1*/FABP1 is downregulated in PGAM5^KO^ HCC and hepatoma cells, likely contributing to reduced fatty acid uptake.^6^ The downregulation of *FASN* suggested that loss of PGAM5 may have a broader impact on lipid metabolism, yet significantly reduced *FASN*/FASN levels (**Figures 3e-g)** and acetyl-CoA carboxylase (ACC), which converts acetyl-CoA to malonyl-CoA in the rate limiting step of fatty acid synthesis, (**Figure 2e**) were only validated in the HepG2 cell line.

### PGAM5 regulates multiple lipid pathways

To determine the global impact of PGAM5 loss on the HCC lipidomes, we performed an unbiased lipidomic analysis in the Huh7 WT and PGAM5^KO^ cells. A threshold of log fold change 1.0 and p-value of 0.05 was used in the analysis. The genotypes distinctly clustered using OPLS-DA, suggesting that the total lipid profile changes considerably in the absence of PGAM5 (**Figure 4a**). The volcano plot from the global analysis of the WT and PGAM5^KO^ cells displays the top upregulated lipids (DG 20:1_22:3, DG 16:1_22:6, DG 18:1_20:3, Cer 20:0;O3/20:0;O) in red color and top downregulated lipids (PS 6:0_6:0, DG 20:0_22:2, TG O-8:0_8:0_8:0) in blue (**Figure 4b)**. A heatmap of lipid species showed the distribution of specific lipids that are upregulated (red) and downregulated (blue) in both WT and PGAM5^KO^ cells (**Figure 4c**). We integrated lipidomic and RNAseq data from PGAM5^KO^ cells to conduct a joint pathway analysis (**Figure 4d**). Fatty acid biosynthesis and degradation, glycerophospholid, glycerolipid, steroid hormone biosynthesis, ether lipid metabolism, and sphingolipid metabolism pathways are enriched in both datasets.

**Figure 4.**
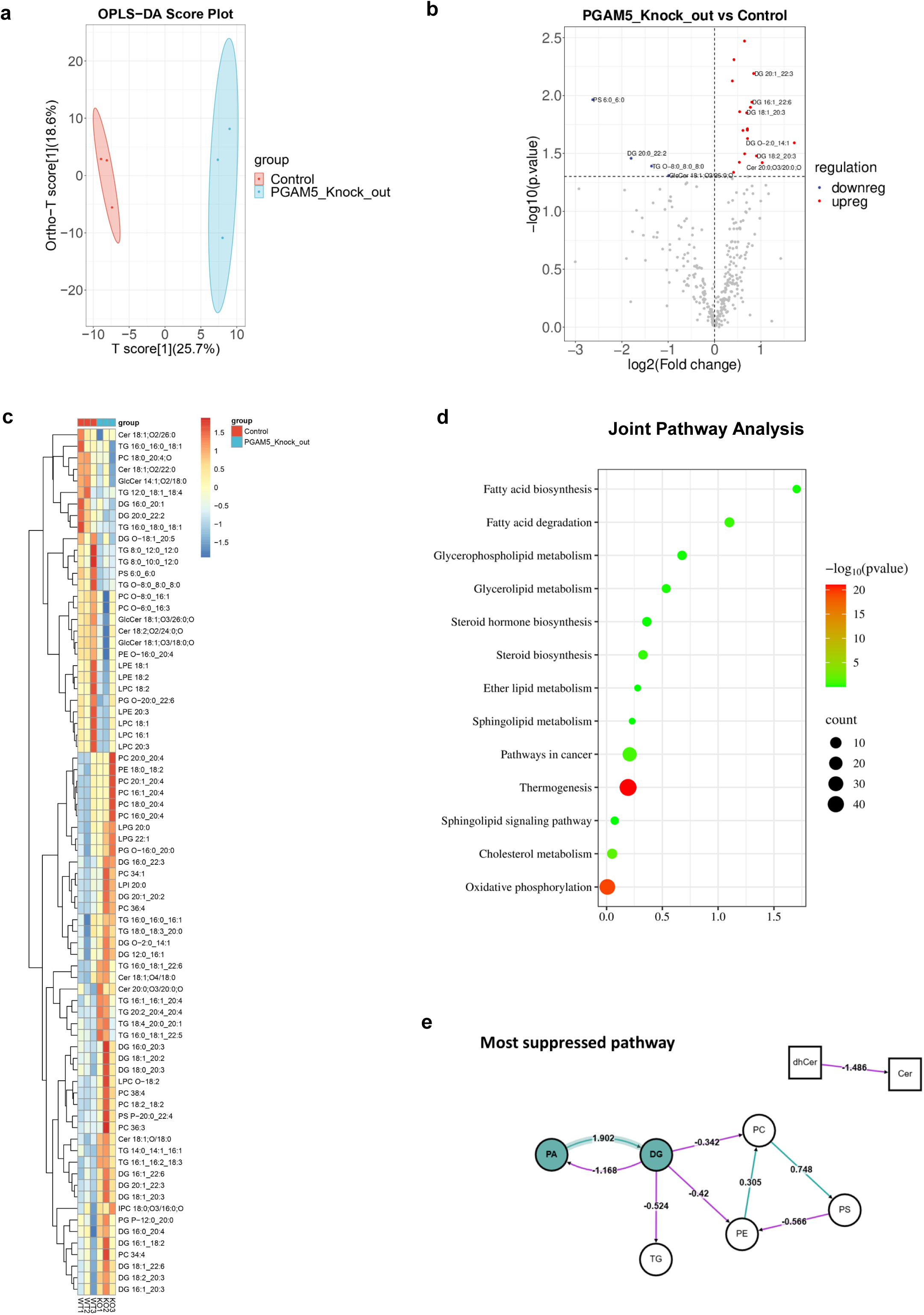
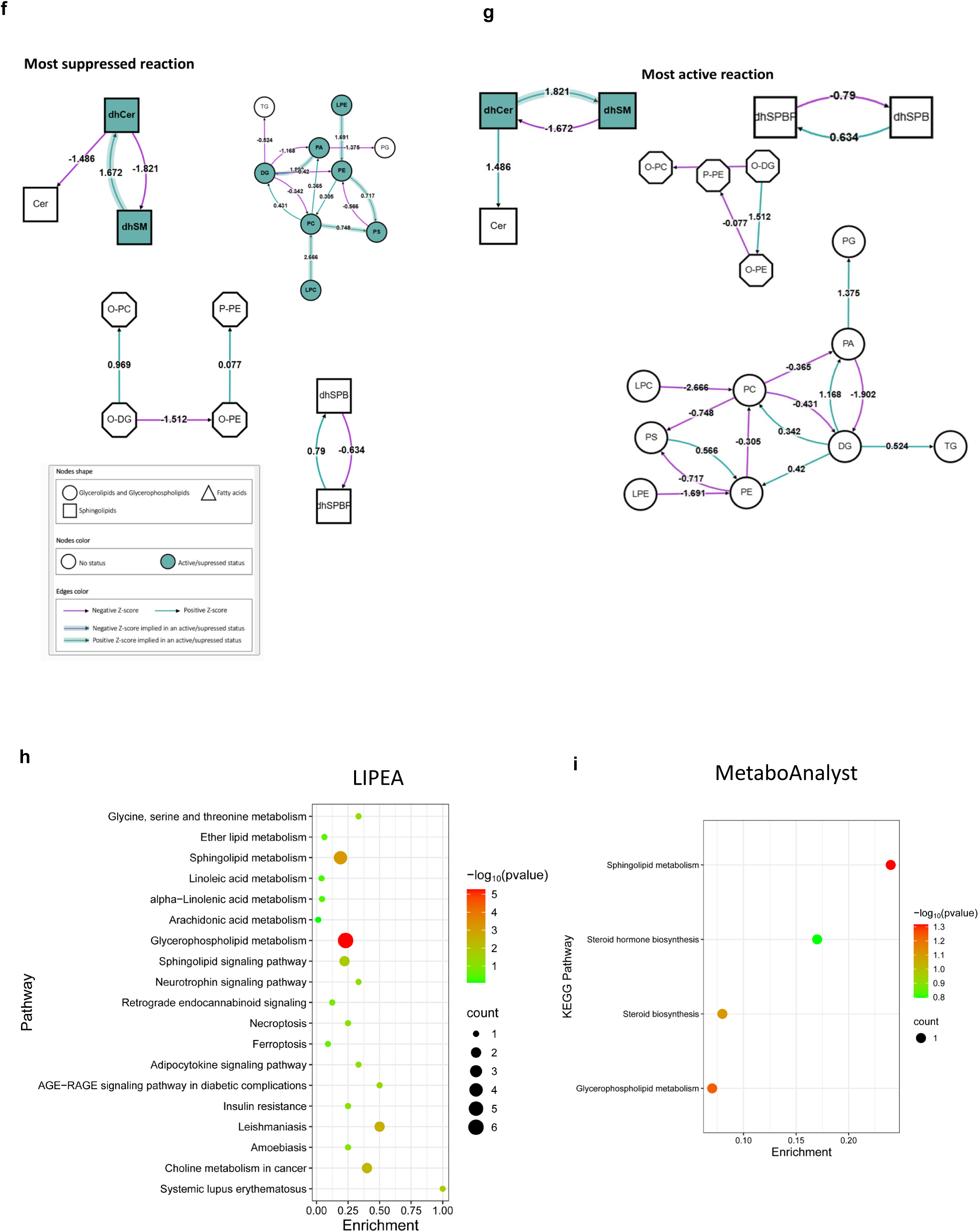
Fatty acid biosynthesis, glycerophospholipid and sphingolipid metabolism are impacted by loss of *PGAM5*. **a.** Orthogonal partial least squares–discriminant analysis (OPLS-DA) distinguishes between WT and PGAM^KO^ Huh7 cells based on lipidomic profiles. The first component (T score [1], 25.7% variance) and the second component (18.6% variance) demonstrate strong group separation. **b.** Differential expression of lipid species between PGAM5^KO^ and WT Huh7 cells. The x-axis represents log2 fold change (PGAM5^KO^ vs. WT), and the y-axis shows –log10 p-values (adjusted). Volcano plot of upregulated (red circles) and downregulated (blue circles) lipid species displayed for PGAM5^KO^ relative to WT in Huh7 cells. **c.** Hierarchical clustering heatmap showing relative abundance of lipid species across replicates of WT and PGAM5^KO^ cells. Rows represent lipid species (grouped by subclass), while columns correspond to biological replicates. The color gradient reflects normalized abundance (Z-scores). Clustering reveals subclass-specific alterations, including significant changes in glycerophospholipids and sphingolipids. **d**. Joint pathway enrichment combining lipidomics and differentially expressed genes (DEGs) from single-cell RNA sequencing of PGAM5^KO^ versus WT Huh7 cells. Circle size corresponds to impact score, and color scale indicates statistical significance (–log10 p-value). **e.** BioPAN pathway analyses of the most suppressed pathways. Lipid class is depicted by shape: Sphingolipids (squares), glycerolipids/ glycerophospholipids (circles), and fatty acids (triangles). Green fill or line highlight identifies a significantly active or suppressed node or pathway. A green edge indicates a positive Z-score and a red edge indicates a negative Z-score; > 1.645 corresponds to p < 0.05. **f.** BioPAN pathway analyses of the most suppressed reactions. **g.** BioPAN pathway analyses of the most active reactions. **h., i.** LIPEA and MetaboAnalyst pathway analysis plots. Circle size corresponds to impact score, and color scale indicates statistical significance (–log10 p-value).

The BioPAN lipid subclass networks for PGAM5^KO^ compared to WT (**Figures 4e-g)**. Pathway analysis revealed a suppression in the conversion of phosphatidic acid (PA) to diacylglycerol (DAG, DG) in the PGAM5^KO^ group (Z-score = 1.902). The most strongly suppressed biosynthetic pathways in the PGAM5^KO^ group were those leading to glycerophospholipids, including phosphatidylserine (PS, Z-score = 2.414), phosphatidylcholine (PC, Z-score = 2.666), and phosphatidylethanolamine (PE, Z-score = 1.691). In contrast, the conversion of dihydroceramide (dhCer) to dihydrosphingomyelin (dhSM) emerged as the most upregulated reaction (Z-score = 1.821; **Table 1**). Dihydroceramides serve as precursors for ceramides, and their de novo synthesis begins in the endoplasmic reticulum with the condensation of L-serine and palmitoyl-CoA.^51^ This process generates dhCer, which is subsequently converted to ceramide by dihydroceramide desaturase (DES). Ceramides then serve as substrates for sphingolipid biosynthesis in the Golgi apparatus. Two isoforms of DES exist, each with distinct tissue expression patterns. Notably, DES1 is widely expressed and also functions as an oxygen sensor, linking ceramide synthesis to the cellular response to hypoxia.^52^ In the PGAM5^KO^ group, reactions involving lysophospholipids as substrates were markedly suppressed, including pathways such as LPC → PC → PS (Z-score = 2.414), LPC → PC → DG (Z-score = 2.190), LPC → PC → PA (Z-score = 2.143), and LPE → PE → PS (Z-score = 1.703). This suppression likely contributes to the accumulation of lysophosphatidylcholine (LPC), which is known to inhibit hepatocellular carcinoma.^53,54^ Additionally, the conversion of diacylglycerol (DG) to triacylglycerol (TG) was also reduced (Z-score = 0.524), consistent with the observed decrease in lipid droplet formation in the PGAM5^KO^ group.^6^ To identify the metabolic pathways affected by *PGAM5* deletion, we performed pathway enrichment analysis with LIPEA and MetaboAnalyst, which showed that the glycerolipid and phospholipid biosynthesis pathways were impacted in *PGAM5^KO^* cells (**Figures 4h, i**).

**Table 1.** Lipid pathway analysis.

| **Lipid class most active reaction (KO vs WT)** |  |  |
| --- | --- | --- |
| **Reactions chains** | **Z-score** | **Predicted genes** |
| dhCer → dhSM | 1.821 | SGMS1, SGMS2 |
| **Lipid class most suppressed reaction (KO vs WT)** |  |  |
| **Reactions chains** | **Z-score** | **Predicted genes** |
| LPC → PC → PS | 2.414 | LPCAT1, LPCAT2, LPCAT3, LPCAT4, PTDSS1 |
| LPC → PC → DG | 2.19 | LPCAT1, LPCAT2, LPCAT3, LPCAT4 |
| LPC → PC → PA | 2.143 | LPCAT1, LPCAT2, LPCAT3, LPCAT4, PLD1, PLD2 |
| PA → DG | 1.902 | PLPP1, PLPP2, PLPP3 |
| LPE → PE → PS | 1.703 | MBOAT1, MBOAT2, LPCAT3, LPCAT4, PTDSS2 |
| dhSM → dhCer | 1.672 | SGMS1, SGMS2, SMPD1 |
| **Lipid class most suppressed pathway (KO vs WT)** |  |  |
| PA → DG | 1.902 | PLPP1, PLPP2, PLPP3 |

### PGAM5 deletion reduces diacylglycerol abundance

Based on the lipidomic data, we derived total abundance of DAG in WT and PGAM5^KO^ Huh7 cells (**Figure 5a**). To validate the reduced DAG concentration, we employed a targeted lipid analysis to quantify cellular DAG concentration. In both the Huh7 and HepG2 models, PGAM5^KO^ cells had significantly lower DAG concentration than WT cells after palmitate treatment (**Figure 5b, c**) implying that PGAM5 regulates PA→ DAG synthesis. Nuclear localization and activation of lipin 1 as a phosphatidic acid phosphatase (PAP) is dictated by its phosphorylation status, which is in part regulated by PGAM5’s phosphatase activity.^55,56^ In the PAP pathway, lipin 1 enzymatically activates conversion of phosphatidic acid (PA) into DAG. Endogenous lipin 1 protein was immunoprecipitated from WT and PGAM5^KO^ Huh7 cells and the phosphorylation state of lipin 1 was determined using PhosTag gel electrophoresis, which tags phosphorylated protein species and slows gel migration (**Figure 5d**). Lipin 1 immunoprecipitated from PGAM5^KO^ cells demonstrated a higher total phosphorylation level than lipin 1 from WT cells, suggesting that loss of PGAM5’s phosphatase activity may be linked to the PAP activity of lipin 1 in HCC (**Figure 5e**). To determine if lipin 1 co-activation dependent gene expression was impacted by *PGAM5* knockout, transcript levels of *FABP1* and *carnitine palmitoyltransferase 1A* (*CPT1a)*, regulator of mitochondrial long chain fatty acid import, were quantified.^56^ Both PGAM5^KO^ cell lines demonstrated significantly reduced transcript levels of *FABP1* and *CPT1a* (**Figure 5f-i**). We hypothesize that the mechanism of PA → DAG synthesis may involve PGAM5’s post-translational modification of lipin 1. Overall, these results demonstrates that *PGAM5* deletion not only modulates mitochondrial homeostasis and bioenergetics, but also impacts glycerolipid and phospholipid metabolic pathways in HCC models.

**Figure 5.**
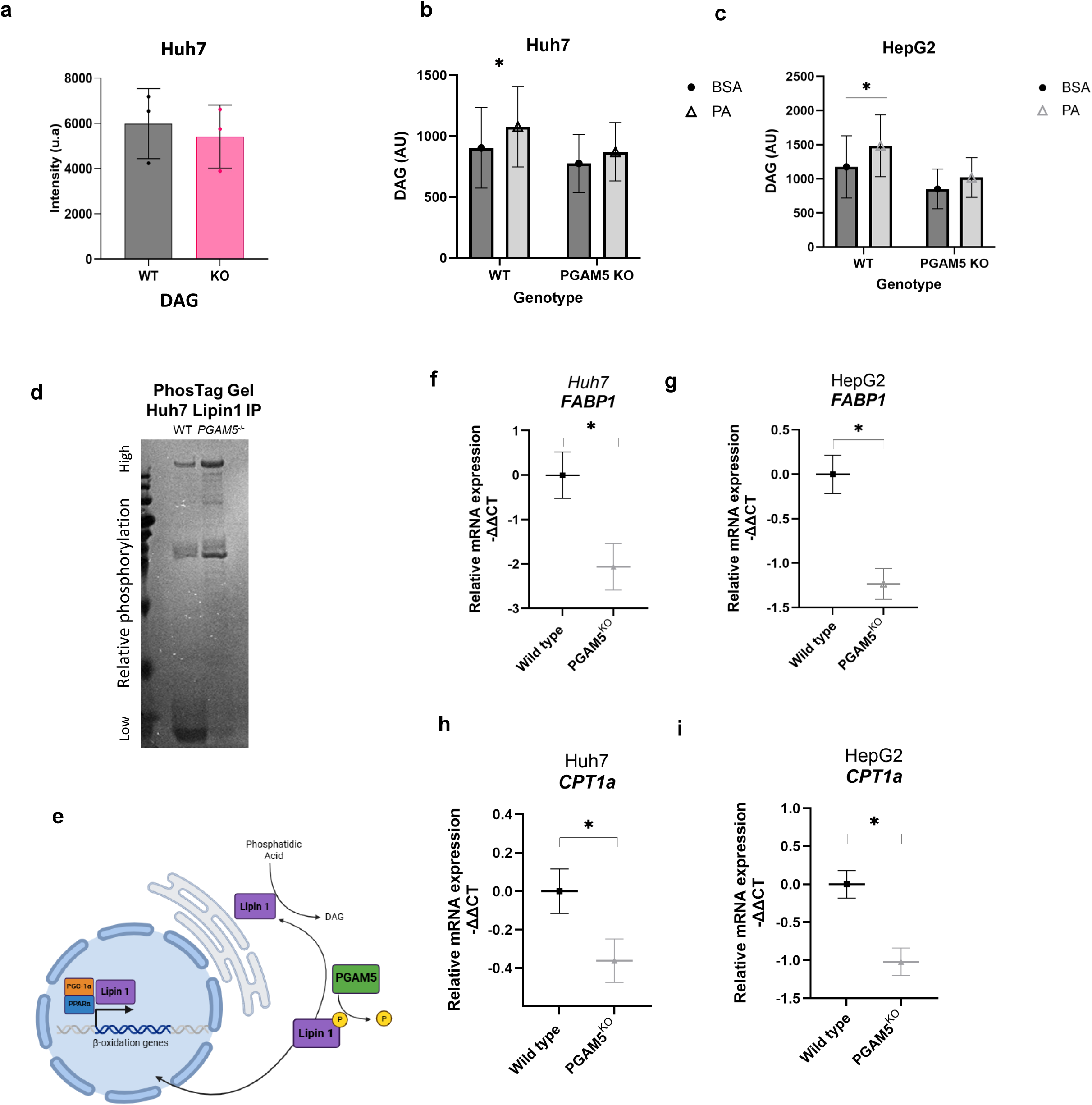
Deletion of PGAM5 reduces cellular diacylglycerol concentration. **a.** Comparative derived total abundance of diacylglycerol (DAG) in the Huh7 lipidome of wildtype (WT) and PGAM5 knockout (KO) cells, mean ± SD. **b. c.** Relative quantification of diacylglycerol (DAG) abundance in wild type (WT) and PGAM5^KO^ HepG2 (f) and Huh7 (g) cells cultured with bovine serum albumin (BSA, filled circles) or palmitic acid (PA, open triangles). AU - arbitrary units. Displayed data are derived from 3 biological replicates with a minimum of 2 technical replicates per sample. **d.** Coomassie blue stained PhosTag gel of lipin-1 protein immunoprecipitated from Huh7 WT and PGAM5 KO Huh7 cells. **e.** Proposed mechanistic link: PGAM5 mediated dephosphorylation of lipin-1 modulates metabolic activity by upregulating DAG synthesis from phosphatidic acid and promoting transcription of genes regulating beta-oxidation. **f.-i.** Real-time (RT) qPCR relative expression of Huh7 (**f, h**) and HepG2 (**g.i**) wild type (black square) and PGAM5 KO (grey triangle) transcripts *fatty acid binding protein 1* (*FABP1*, **f, g**) and *carnitine palmitoyltransferase 1A* (*CPT1a*, **h, i**). RT qPCR data are derived from a minimum of 3 biological replicates with 2 technical replicates per sample, mean ± SEM; *, p<.05.

## Discussion

Excess dietary lipid intake is initially associated with increases in hepatic mitochondrial mass and upregulation of mitochondrial oxidative function, encompassing changes in beta-oxidation, hepatic TCA cycle, ketogenesis, respiratory chain activity, and ATP synthesis.^57,58^ As lipid accumulates in the liver, these adaptive metabolic mechanisms transition to a maladaptive phenotype, wherein mitochondria have incomplete beta-oxidation, toxic intermediates accumulate, ketogenesis is impaired, and respiration, albeit with maintenance of TCA cycle function.^58,59^ These bioenergetic changes are progressive and in a subset of patients this fosters the transition to steatohepatitis and carcinoma.^60–62^ Based on compiled lipidomic analyses of human HCCs, short chain fatty acids, monounsaturated fatty acids, and sphingomyelins are increased in HCC; whereas, polyunsaturated fatty acids are typically decreased.^59,63,64^ Despite the increased attention on the HCC lipidome over recent years, how the milieu of intracellular lipid species impact cancer progression and therapy remains unclear.

PGAM5 expression in HCC is correlated with reduced overall survival in human patients.^10,65^ Upregulation of PGAM5 may confer a selective advantage in tumors through preservation of mitochondrial function under oxidative and metabolic stress.^27,28,66^ It is well established that PGAM5 mediates mitochondrial homeostasis by regulating biogenesis, fission, and mitophagy.^7,8,26,43^ Loss of PGAM5 has been previously shown to result in accumulation of damaged mitochondria; we have corroborated this finding in HCC cell models and correlated it to abnormal macronutrient metabolism.^41,45,48^ Ultimately, downregulation of mitochondrial respiration reduced energetic bioavailability, which was further exacerbated by diminished lipid droplet stores. These data provide a mechanistic link between mitochondrial dysfunction and remodeling of metabolic pathways in cancer.

This work is the first to characterize the transcriptional and lipidomic landscape associated with *PGAM5* deletion in HCC.^8,27,28,67^ We have shown that deletion of *PGAM5* downregulates transcription of lipid metabolic mediators and nuclear-encoded electron transport cycle genes, and promotes energetic reliance on non-lipid substrates. Yet rather than solely dysregulating beta oxidation, knockout of *PGAM5* specifically limits long chain fatty import, which supports a model of coordinated suppression of lipid uptake and utilization. Further, we have shown that the glycerophospholipid and lysophospholipid pathways are suppressed in PGAM5^KO^ cells, resulting in reduced synthesis of DAG and subsequent accumulation of LPC, likely worsening ROS induced mitochondrial injury.^68,69^

Studies from our group and others indicate that suppression of PGAM5 expression enhances cellular susceptibility to injury.^21,28,32,65^ Individually and collectively, these findings reveal vulnerabilities in metabolic adaptability that may be therapeutically exploited in HCC. The conserved overexpression of PGAM5 across species further supports its potential as a therapeutic target. Together, these results raise important mechanistic questions regarding the contribution of specific lipid species to tumor progression and the potential utility of phosphatase inhibition as a therapeutic strategy in metabolic disease and HCC. Further detailed pathway analyses are required to fully characterize PGAM5’s network in lipid metabolism and cancer.

## Acknowledgments

The authors acknowledge the technical assistance provided by the GeneLab Center for Experimental Infectious Disease Research and Clinical Pathology Laboratory at the Louisiana State University School of Veterinary Medicine and the Histology Laboratory at the Louisiana Animal Disease Diagnostic Laboratory. The authors would also like to acknowledge IDeA National Resource for Quantitative Proteomics and Arkansas Children’s Research Institute for RNA sequencing bioinformatics analysis, Fabrizio Donnarumma at the Louisiana State University Mass Spectrometry Facility for Lipidomics Sample Processing, Alicia Withrow at the Michigan State University Center for Advanced Microscopy for EM sample preparation and Dalen Agnew at Michigan State University for EM image acquisition.

## Data availability

The lipidomic and RNA sequencing datasets are available at (https://doi.org/10.5061/dryad.gtht76j0x). The other datasets generated during and/or analyzed during the current study are available from the corresponding author upon reasonable request.

## Funding and additional information

This research was funded by the National Institutes of Health under Award Number 1P20GM135000-01A1 (A.J.). M.R.G. was supported by the National Institute of General Medical Sciences of the National Institutes of Health under award number R35GM150564.

## Author contributions

A.N.J. and M.R.G. Conceptualization; P.K.G., G.M., P.D., D.A., H.G., and T.B. Data curation; M.R.G., P.K.G., P.K.G., C.-C.L., H.G., J.M., and A.N.J. Formal analysis; M.R.G. and A.N.J. Funding acquisition; M.R.G., P.K.G., G.M., P.D., E.S., D.A., and A.J. Investigation; M.R.G., P.K.G., G.M., and A.N.J. Methodology; M.R.G., H.G., J.M., D.A., and A.N.J. Project administration; M.R.G. and A.N.J. Resources; P.K.G., G.M., J.M., and C.-C.L. Software; M.R.G. and A.N.J. Supervision; M.R.G., P.K.G., G.M., P.D., C.-C.L., and A.J. Validation; M.R.G., P.K.G., G.M., P.D., C.-C.L., and A.J. Visualization; M.R.G., P.K.G., and A.N.J. Roles/Writing - original draft; M.R.G., P.K.G., G.M., P.D., C.-C.L., E.S., H.G., J.M., D.A., T.B., and A.N.J. Writing - review & editing.

